## Supplementary Table S2 for "Passive accumulation of alkaloids in inconspicuously colored frogs refines the evolutionary paradigm of acquired chemical defenses"

**Table S1**. A summary of data available on alkaloid detection in “undefended” lineages of poison frogs prior to this study.

| **Species** | **Method** | **Reported Results** | **Our interpretation** | **Rationale if our interpretation is distinct from reported results** | **Reference** |
| --- | --- | --- | --- | --- | --- |
| *Allobates femoralis* | GC-MS | trace amount of a 5,8-indolizidine alkaloid detected in one individual from one of six tested populations | low levels of an alkaloid present in one skin, possibly in others below detection threshold | N/A | [1] |
| *Allobates femoralis* | TLC | no spots detected in 15 individuals | no alkaloids detected, but image of TLC plate not available to verify interpretation | N/A | [2] |
| *Allobates femoralis* | mouse bioassay and predator assay | no difference in mouse behavior between saline and *A. femoralis* injection, and naïve chicks readily ate three frogs | skin secretion does not contain [enough] compounds that irritate mice or deter chicks | N/A | [3] |
| *Allobates femoralis* | mouse bioassay | difference in mouse behavior between saline and *A. femoralis* injection | skin extracts contain compounds that irritate mice, but unclear if those compounds are lipophilic alkaloids, and others have speculated that irritant effects are due to the benzocaine used for euthanasia [4] | N/A | [5] |
| *Allobates femoralis* | GC-MS | no alkaloids detected in two individuals | no alkaloids detected, but we cannot rule out the occurrence of lipophilic alkaloids below detection thresholds or not in the Daly database | N/A | [4] |
| *Allobates femoralis* | GC-MS | no alkaloids detected in one individual | no alkaloids detected, but we cannot rule out the occurrence of lipophilic alkaloids below detection thresholds or not in the Daly database | the study tested ethanol in which whole animals were soaked after being collected | [6] |
| *Allobates femoralis* | feeding assay (alkaloid-dusted fruit flies) followed by GC-MS | a captive-bred individual had detectable sparteine in skin after a month | the single captive-bred individual tested accumulated provisioned sparteine | N/A | [7] |
| *Allobates femoralis* | oral administration assay (alkaloids dissolved in 50% EtOH) followed by GC-MS | six wild-caught frogs had DHQ in trace quantities (~1% of amount orally administered) in skins, livers, and feces (three individuals), but no HTX **235A** (three individuals) after 14 days | the frogs accumulate DHQ but not HTX **235A** | N/A | [8] |
| *Allobates insperatus* | TLC | no spots detected in 12 individuals | no alkaloids detected, but image of TLC plate not available to verify interpretation | N/A | [2] |
| *Allobates kingsburyi* | TLC | spot observable in three individuals | spot on TLC plate suggests presence of skin metabolites (potentially alkaloids) | the negative control used in this experiment (*H. yasuni*) yielded spots on the TLC plate but was assumed to lack alkaloids, which likely confounded the interpretation of results | [9] |
| *Allobates myersi* | GC-MS | no alkaloids detected in one individual | no alkaloids detected, but we cannot rule out the occurrence of lipophilic alkaloids below detection threshold | N/A | [1] |
| *Allobates sumtuosus* | GC-MS | no alkaloids detected in two individuals | no alkaloids detected, but we cannot rule out the occurrence of lipophilic alkaloids below detection thresholds or not in the Daly database | the study tested ethanol in which whole animals were soaked after being collected | [6] |
| *Allobates talamancae* | either by mouse bioassay or [^3^H]saxitoxin binding assay | no alkaloids detected (likely in one individual) | no evidence of lipophilic alkaloid presence or absence | water-based extraction process would not retain high levels of lipophilic alkaloids, so cannot be used as evidence of presence or absence | [10,11] |
| *Allobates talamancae* | GC-MS | no alkaloids detected in 10 individuals | no alkaloids detected, but we cannot rule out the occurrence of lipophilic alkaloids below detection thresholds or not in the Daly database | the study tested ethanol in which whole animals were soaked after being collected | [6] |
| *Allobates talamancae* | feeding assay (alkaloid-dusted fruit flies) followed by GC-MS | no alkaloids detected in unknown number of individuals | no evidence for alkaloid accumulation after five weeks | N/A | [12] |
| *Allobates talamancae* | unknown (possibly GC-MS) | no alkaloids detected in unknown number of wild-caught individuals | no alkaloids detected, but we cannot rule out the occurrence of lipophilic alkaloids below detection threshold | N/A | unpub. data reported in [13] |
| *Allobates zaparo* | TLC | either no or weak (at most) spots detected in 20 individuals | spot on TLC plate suggests presence of skin metabolites (potentially alkaloids) | Darst et al. [2] bin this species into a non-toxic category despite presence of light spots | [2] |
| *Allobates zaparo* | mouse bioassay and predator assay | no difference in mouse behavior between saline and *A. zaparo* injection, and naïve chicks readily ate six frogs | skin secretion does not contain [enough] compounds that irritate mice or deter chicks | N/A | [14] |
| *Allobates zaparo* | mouse bioassay | no difference in mouse behavior between saline and *A. zaparo* injection, using skin samples from five individuals | skin secretion does not contain [enough] compounds that irritate mice | N/A | [3] |
| *Aromobates nocturnus* | TLC, GC-MS, and thermospray mass spectrometry, and subcutaneous injection into white mice, all using methanol extract | no alkaloids detected and no effect on mice in extracts from 10 adult females | no alkaloids detected nor enough compounds that irritate mice, but we cannot rule out the occurrence of lipophilic alkaloids below detection threshold | N/A | [15] |
| *Aromobates nocturnus* | [^3^H]saxitoxin binding assay | no TTX detected in unknown number of individuals | unknown if lipophilic alkaloids present or absent | water-based extraction process would not retain high levels of lipophilic alkaloids, so cannot be used as evidence of presence or absence | [11] |
| *Colostethus imbricolus* | mouse bioassay and [^3^H]saxitoxin binding assay | no TTX detected in saxitoxin assay, but very marginal effects were noted in mouse assay, based on one skin sample | no direct evidence of lipophilic alkaloids, but potentially present | water-based extraction process would not retain high levels of lipophilic alkaloids, so cannot be used as evidence of presence or absence | [11,15] |
| *Colostethus imbricolus* | unknown (possibly GC-MS) | no lipophilic alkaloids detected in one skin sample | no alkaloids detected, but we cannot rule out the occurrence of lipophilic alkaloids below detection threshold | N/A | [15] |
| *Colostethus panamansis* (identified as “Colostethus inguinalis” in [11]) | feeding assay (alkaloid-dusted fruit flies) followed by GC-MS | no alkaloids detected in unknown number of individuals | no evidence for alkaloid accumulation after five weeks | N/A | [12] |
| *Colostethus panamansis* (identified as “Colostethus inguinalis” in [11]) | mouse bioassay, [^3^H]saxitoxin binding assay, HPLC, and TLC | TTX detected in three individuals | unknown if lipophilic alkaloids present or absent | water-based extraction process would not retain high levels of lipophilic alkaloids, so cannot be used as evidence of presence or absence | [11] |
| “*Colostethus* sp.” (identified as “Colostethus new species”, sympatric with *Aromobates nocturnus*, Trujillo, Venezuela in [11]) | either by mouse bioassay or by [^3^H]saxitoxin binding assay | no alkaloids detected (likely in one individual) | unknown if lipophilic alkaloids present or absent | water-based extraction process would not retain high levels of lipophilic alkaloids, so cannot be used as evidence of presence or absence | [11] |
| *Colostethus ucumari* | methanol extract (i.e., lipophilic compounds) caused minor agitation and twitching in mice, but aqueous extract (i.e., water-soluble compounds) caused gagging and some inactivity | non-toxic methanol extract (lipophilic compounds), non-toxic, aqueous extract (hydrophilic compounds), both from one individual (AMNH 104371) | evidence for presence of irritating lipophilic and hydrophilic compounds (possibly alkaloids) | N/A | [16] |
| *Colostethus ucumari* | GC-MS of methanol extract | small amounts of multiple lipophilic compounds not corresponding to known dendrobatid alkaloids from one individual (AMNH 104371) | lipophilic compounds present, potentially alkaloids | N/A | [16] |
| *Colostethus ucumari* | high-resolution GC-MS of methanol extract | only trace amount of one of lipophilic compounds detectable from 30-year-old methanol extract (AMNH 104371), which turned out to be artifact (2-benzothazolyl-N,N-dimethyl dithiocarbamate) | identity of lipophilic compounds remains unknown | N/A | [16] |
| *Ectopoglossus saxatilis* | GC-MS of methanol extract | no alkaloids detected in one individual | no alkaloids detected, but we cannot rule out the occurrence of lipophilic alkaloids below detection threshold | N/A | unpub. data reported in [17] |
| *Hyloxalus awa* | predator assay | naïve chicks readily ate unknown number of frogs | animal does not contain [enough] compounds that deter predators | this species was used as a control in predation experiments | [14] |
| *Hyloxalus azureiventris* | oral administration assay followed by GC-MS, feeding assay (alkaloid dusted fruit flies) followed by GC-MS | species sequestered four alkaloids dissolved in methanol-saline solution, but later studies found no sequestered alkaloids from dusted fruit flies, in unknown number of individuals | could sequester measurable levels of alkaloids, though likely inefficiently given the conflict between feeding and oral administration assays | N/A | unpub. data as reported in  [18] |
| *Hyloxalus azureiventris* | unknown (possibly GC-MS) | no alkaloids detected in unknown number of wild-caught individuals | no alkaloids detected, but we cannot rule out the occurrence of lipophilic alkaloids below detection threshold | N/A | unpub. data reported in [13] |
| *Hyloxalus chlorocraspedus* | unknown (possibly GC-MS) | no alkaloids detected in unknown number of individuals | no alkaloids detected, but we cannot rule out the occurrence of lipophilic alkaloids below detection threshold | N/A | unpub. data as reported in [19] |
| *Hyloxalus elachyhistus*  (identified as “*Hyloxalus infraguttatus*” in [12]) | either by mouse bioassay or by [^3^H]saxitoxin binding assay | no alkaloids detected (likely) in one individual | unknown if lipophilic alkaloids present or absent | water-based extraction process would not retain high levels of lipophilic alkaloids, so cannot be used as evidence of presence or absence | [11] |
| *Hyloxalus elachyhistus* (identified as “*Hyloxalus infraguttatus*” in [20]) | targeted GC-MS with nicotine standard followed by filtering against database of poison-frog alkaloids [21] | trace levels of diverse alkaloids, mostly indolizidines and histrionicotoxins, detected in study of nine individuals | presence of skin alkaloids | N/A | [20] |
| *Hyloxalus jhoncito* | GC-MS of methanol extract | no alkaloids detected in one individual | no alkaloids detected, but we cannot rule out the occurrence of lipophilic alkaloids below detection thresholds or not in the Daly database | N/A | unpub. data reported in [22] |
| *Hyloxalus nexipus* | TLC | spot observable in one of three individuals | spot on TLC plate suggests presence of skin metabolites (potentially alkaloids) | the negative control used in this experiment (*H. yasuni*) yielded a spot on the TLC plate but was assumed to lack alkaloids, which likely confounded the interpretation of results | [9] |
| *Hyloxalus sauli* | TLC | no spots detected in 10 individuals | no alkaloids detected, but image of TLC plate not available to verify interpretation | N/A | [2] |
| *Hyloxalus yasuni* (identified as “*Colostethus* sp. D” in [2]) | TLC | no spots detected in 22 individuals | no alkaloids detected, but image of TLC plate not available to verify interpretation | N/A | [2] |
| *Hyloxalus yasuni* (called “*Hyloxalus maculosus*” in [9]) | TLC | used as negative control, but spots were observable in both individuals (one spot in one individual, at least three in the other) | spots on TLC plate suggest presence of skin metabolites (potentially alkaloids) | used as negative control in [9] due to earlier TLC result for [2]) | [9] |
| *Hyloxalus vertebralis* | TLC | spot observable in two of five individuals | spots on TLC plate suggest presence of skin metabolites (potentially alkaloids) | the negative control used in this experiment (*H. yasuni*) yielded spots on the TLC plate but was assumed to lack alkaloids, which likely confounded the interpretation of results | [9] |
| *Leucostethus fraterdanieli*  (identified as “*Colostethus* sp. common nr Villa Maria, Caldas, Colombia in [11]) | either by mouse bioassay or by [^3^H]saxitoxin binding assay | no alkaloids detected (likely in one individual) | unknown if lipophilic alkaloids present or absent | water-based extraction process would not retain high levels of lipophilic alkaloids, so cannot be used as evidence of presence or absence | [11] |
| *Leucostethus fugax* | TLC | two spots observable in four individuals (one dark and one faint) and one spot observable in one individual | spots on TLC plate suggest presence of skin metabolites (potentially alkaloids) | the negative control used in this experiment (*H. yasuni*) yielded spots on the TLC plate but was assumed to lack alkaloids, which likely confounded the interpretation of results | [9] |
| *Leucostethus siapida* | GC-MS of methanol extract | no alkaloids detected in one individual | no alkaloids detected, but we cannot rule out the occurrence of lipophilic alkaloids below detection thresholds or not in the Daly database | N/A | unpub. data reported in [23] |
| *Mannophryne* cf. *collaris* (identified as “*Colostethus* new species, cf. *collaris*” Trujillo, Venezuela in [12]) | either by mouse bioassay or [^3^H]saxitoxin binding assay | no alkaloids detected (likely in one individual) | unknown if lipophilic alkaloids present or absent | water-based extraction process would not retain high levels of lipophilic alkaloids, so cannot be used as evidence of presence or absence | [11] |
| *Mannophryne riveroi* | either by mouse bioassay or by [^3^H]saxitoxin binding assay | no alkaloids detected (likely in one individual) | unknown if lipophilic alkaloids present or absent | water-based extraction process would not retain high levels of lipophilic alkaloids, so cannot be used as evidence of presence or absence | [11] |
| *Mannophryne trinitatis* | either by mouse bioassay or by [^3^H]saxitoxin binding assay | no alkaloids detected (likely in one individual) | unknown if lipophilic alkaloids present or absent | water-based extraction process would not retain high levels of lipophilic alkaloids, so cannot be used as evidence of presence or absence | [11] |
| *Paruwrobates erythromos* | GC-MS | three alkaloids detected across five individuals | presence of skin alkaloids | N/A | [1,24] |
| *Silverstoneia flotator* | 14-mo avian diet (regurgitate) study and herpetofauna census | 91 sympatric bird species do not consume *S. flotator* despite its being one of the two most common anurans | indirect evidence that animals may have compounds (possibly alkaloids) that deter predators | N/A | [25] |
| *Silverstoneia flotator* | feeding experiment with bat species (*Trachops cirrhosus*) | *Trachops cirrhosus* bats (from Barro Colorado Island, Panama) spit out unknown number of *S. flotator* individuals from Peninsula Gigante population (Panama) when hand-fed the frogs (field seasons 1979–81), and one bat even died as a result of frog consumption; it is highly unusual for this bat species to die after predation (Michael J. Ryan, pers. comm.) | indirect evidence that animals may have compounds (possibly alkaloids) that deter predators |  | (Michael J. Ryan, pers. comm.) |
| *Silverstoneia flotator* | GC-MS | no alkaloids detected in one individual | no alkaloids detected, but we cannot rule out the occurrence of lipophilic alkaloids below detection thresholds or not in the Daly database | the study tested ethanol in which whole animals were soaked after being collected | [6] |
| *Silverstoneia flotator* | unknown (possibly GC-MS) | no alkaloids detected in unknown number of individuals | no alkaloids detected, but we cannot rule out the occurrence of lipophilic alkaloids below detection threshold | N/A | Daly, unpub. data as reported in [6,25] |
| *Silverstoneia punctiventris* | head-space solid phase microextraction coupled to GC-MS (targeted method) | 13 compounds with alkaloid structures detected (including six from Daly’s poison-frog alkaloid database) across eight individuals | presence of six lipophilic alkaloids previously documented in poison frogs, seven other alkaloids | N/A | [26] |
