## Supplementary Table S3 for "Passive accumulation of alkaloids in inconspicuously colored frogs refines the evolutionary paradigm of acquired chemical defenses"

**Table S3.** Collection localities, specimen numbers, size, sex, and summary of alkaloid quantities and diversity for each individual. ND, not determined; F, female; M, male; J, juvenile.

| **Family** | **Species** | **Total Integrated Area / 1x10^6^** | **Total Number Alkaloids** | **Locality, Province/Department, Country:**  **Latitude, Longitude** | **Preparation Day** | **Museum Number** | **Field Number** | **SVL (mm)**  **mass (g)** | **Sex** |
| --- | --- | --- | --- | --- | --- | --- | --- | --- | --- |
| Bufonidae | *Amazophrynella siona* | 1.350 | 1 | Canelos, Pastaza, EC:  -1.586748, -77.772904 | 08 Oct 2014 | QCAZ:A:58313 | RDT0093 | 15.52  0.4 | ND |
| Bufonidae | *Amazophrynella siona* | 1.795 | 1 | Canelos, Pastaza, EC:  -1.586748, -77.772904 | 08 Oct 2014 | QCAZ:A:58314 | RDT0094 | 16.98  0.5 | ND |
| Bufonidae | *Atelopus* aff*. spurrelli* | 0.107 | 4 | Termales, Chocó, CO:  5.608327 -77.431684 | 17 Nov 2014 | ANDES:A:2438 | RDT0130 | 22.69  0.7 | ND |
| Dendrobatidae | *Allobates insperatus* | 4.697 | 5 | Laguna de San Pedro, Orellana, EC:  -0.241134, -76.946461 | 15 Oct 2014 | QCAZ:A:58323 | RDT0103 | 16.13  0.4 | ND |
| Dendrobatidae | *Allobates insperatus* | 3.519 | 7 | Laguna de San Pedro, Orellana, EC:  -0.241134, -76.946461 | 15 Oct 2014 | QCAZ:A:58324 | RDT0104 | 18.45  0.5 | F |
| Dendrobatidae | *Allobates insperatus* | 2.958 | 4 | Laguna de San Pedro, Orellana, EC:  -0.241134, -76.946461 | 15 Oct 2014 | QCAZ:A:58325 | RDT0105 | 15.73  0.4 | ND |
| Dendrobatidae | *Allobates insperatus* | 0.706 | 1 | Laguna de San Pedro, Orellana, EC:  -0.241134, -76.946461 | 15 Oct 2014 | QCAZ:A:58326 | RDT0106 | 16.67  0.5 | F |
| Dendrobatidae | *Allobates insperatus* | 5.026 | 9 | Laguna de San Pedro, Orellana, EC:  -0.240427, -76.947757 | 15 Oct 2014 | QCAZ:A:58331 | RDT0111 | 17.18  0.5 | F |
| Dendrobatidae | *Allobates insperatus* | 5.076 | 6 | Laguna de San Pedro, Orellana, EC:  -0.240427, -76.947757 | 15 Oct 2014 | QCAZ:A:58332 | RDT0112 | 16.95  0.4 | ND |
| Dendrobatidae | *Allobates insperatus* | 2.338 | 4 | Laguna de San Pedro, Orellana, EC:  -0.240427, -76.947757 | 16 Oct 2014 | QCAZ:A:58333 | RDT0113 | 16.09  0.4 | ND |
| Dendrobatidae | *Allobates insperatus* | 2.036 | 5 | Laguna de San Pedro, Orellana, EC:  -0.240427, -76.947757 | 16 Oct 2014 | QCAZ:A:58334 | RDT0114 | 16.42  0.4 | ND |
| Dendrobatidae | *Allobates juanii* | 1.330 | 1 | Santa Maria, Boyacá, CO:  4.847583, -73.271048 | 03 Nov 2014 | ANDES:A:2425 | RDT0117 | 14.48  0.8 | ND |
| Dendrobatidae | *Allobates kingsburyi* | 0.830 | 2 | Zumbi, Zamora-Chinchipe, EC:  -3.889233, -78.78612 | 28 Sep 2014 | QCAZ:A:58288 | RDT0068 | 17.35  0.4 | ND |
| Dendrobatidae | *Allobates talamancae* | 11.682 | 4 | Salero, Chocó, CO:  5.330797, -76.629868 | 20 Nov 2014 | ANDES:A:2448 | RDT0140 | 25.3  1.6 | F |
| Dendrobatidae | *Allobates talamancae* | 2.929 | 3 | Salero, Chocó, CO:  5.330797, -76.629868 | 20 Nov 2014 | ANDES:A:2449 | RDT0141 | 23.27  1.05 | F |
| Dendrobatidae | *Allobates talamancae* | 3.580 | 2 | Salero, Chocó, CO:  5.330797, -76.629868 | 20 Nov 2014 | ANDES:A:2450 | RDT0142 | 24.62  1.25 | F |
| Dendrobatidae | *Allobates zaparo* | 19.363 | 8 | near Jatún Sacha, Napo, EC:  -1.06509, -77.616818 | 13 Oct 2014 | QCAZ:A:58322 | RDT0102 | 27.94  1.8 | ND |
| Dendrobatidae | *Ameerega bilinguis* | 3,482.601 | 133 | near Jatún Sacha, Napo, EC:  -1.065357, -77.616868 | 12 Oct 2014 | QCAZ:A:58321 | RDT0101 | 20.64  0.9 | ND |
| Dendrobatidae | *Ameerega hahneli* | 596.492 | 85 | Canelos, Pastaza, EC:  -1.584396, -77.789511 | 08 Oct 2014 | QCAZ:A:58309 | RDT0089 | 21.86  1.0 | ND |
| Dendrobatidae | *Ameerega hahneli* | 4,813.290 | 132 | Canelos, Pastaza, EC:  -1.584396, -77.789511 | 08 Oct 2014 | QCAZ:A:58310 | RDT0090 | 24.57  1.1 | F |
| Dendrobatidae | *Ameerega hahneli* | 2,301.534 | 140 | Laguna de San Pedro, Orellana, EC:  -0.240408, -76.947668 | 15 Oct 2014 | QCAZ:A:58327 | RDT0107 | 18.88  0.5 | ND |
| Dendrobatidae | *Ameerega hahneli* | 2,921.383 | 125 | Laguna de San Pedro, Orellana, EC:  -0.240408, -76.947668 | 15 Oct 2014 | QCAZ:A:58328 | RDT0108 | 18.80  0.5 | ND |
| Dendrobatidae | *Andinobates fulguritus* | 533.092 | 85 | Termales, Chocó, CO:  5.611517, -77.418186 | 17 Nov 2014 | ANDES:A:2436 | RDT0128 | 14.60  0.3 | M |
| Dendrobatidae | *Andinobates fulguritus* | 804.436 | 80 | Termales, Chocó, CO:  5.611517, -77.418186 | 17 Nov 2014 | ANDES:A:2437 | RDT0129 | 14.83  0.3 | F |
| Dendrobatidae | *Andinobates minutus* | 142.306 | 73 | La Barra, Valle del Cauca, CO:  3.985532, -77.376676 | 26 Nov 2014 | ANDES:A:2451 | RDT0143 | 13.29  0.3 | ND |
| Dendrobatidae | *Andinobates minutus* | 126.729 | 80 | La Barra, Valle del Cauca, CO:  3.985532, -77.376676 | 26 Nov 2014 | ANDES:A:2452 | RDT0144 | 11.76  0.2 | J |
| Dendrobatidae | *Andinobates minutus* | 38.930 | 59 | La Barra, Valle del Cauca, CO:  3.985532, -77.376676 | 26 Nov 2014 | ANDES:A:2453 | RDT0145 | 11.23  0.2 | J |
| Dendrobatidae | *Andinobates minutus* | 15.724 | 34 | La Barra, Valle del Cauca, CO:  3.985532, -77.376676 | 26 Nov 2014 | ANDES:A:2454 | RDT0146 | 11.73  0.2 | J |
| Dendrobatidae | *Dendrobates truncatus* | 512.289 | 111 | Melgar, Tolima, CO:  4.19277, -74.630953 | 07 Nov 2014 | ANDES:A:2427 | RDT0119 | 20.58  0.8 | M |
| Dendrobatidae | *Dendrobates truncatus* | 738.133 | 115 | Melgar, Tolima, CO:  4.19277, -74.630953 | 07 Nov 2014 | ANDES:A:2428 | RDT0120 | 20.13  0.7 | M |
| Dendrobatidae | *Dendrobates truncatus* | 25,258.195 | 172 | Melgar, Tolima, CO:  4.19277, -74.630953 | 07 Nov 2014 | ANDES:A:2429 | RDT0121 | 24.88  1.1 | F |
| Dendrobatidae | *Epipedobates* aff. *espinosai* | 593.100 | 131 | Cube, Esmeraldas, EC:  0.409176, -79.674679 | 14 Sep 2014 | QCAZ:A:58234 | RDT0014 | 18.00  0.6 | F |
| Dendrobatidae | *Epipedobates* aff. *espinosai* | 101.968 | 83 | Valle Hermoso, Santo Domingo, EC:  -0.085308, -79.273473 | 16 Sep 2014 | QCAZ:A:58249 | RDT0029 | 16.47  0.3 | ND |
| Dendrobatidae | *Epipedobates espinosai* | 1,837.035 | 146 | Río Palenque, Santo Domingo, EC:  -0.590298, -79.362069 | 18 Sep 2014 | QCAZ:A:58270 | RDT0050 | 16.52  0.5 | F |
| Dendrobatidae | *Epipedobates espinosai* | 149.686 | 85 | Mindo, Pichincha, EC:  -0.074375, -78.754373 | 24 Sep 2014 | QCAZ:A:58285 | RDT0065 | 18.00  0.5 | ND |
| Dendrobatidae | *Epipedobates anthonyi* | 832.323 | 127 | Moromoro, El Oro, EC:  -3.65462, -79.741854 | 29 Sep 2014 | QCAZ:A:58290 | RDT0070 | 17.98  0.9 | ND |
| Dendrobatidae | *Epipedobates machalilla* | 6.361 | 38 | Río Ayampe, Manabí, EC:  -1.679337, -80.798273 | 01 Oct 2014 | QCAZ:A:58296 | RDT0076 | 15.36  0.3 | ND |
| Dendrobatidae | *Epipedobates machalilla* | 0.435 | 8 | Río Ayampe, Manabí, EC:  -1.679534, -80.798372 | 01 Oct 2014 | QCAZ:A:58298 | RDT0078 | 14.62  0.3 | ND |
| Dendrobatidae | *Epipedobates tricolor* | 191.854 | 114 | San José de Tambo, Bolívar, EC:  -1.941498, -79.170677 | 04 Oct 2014 | QCAZ:A:58302 | RDT0082 | 21.60  0.8 | ND |
| Dendrobatidae | *Epipedobates tricolor* | 94.105 | 91 | Guanujo, Bolívar, EC:  -1.480303, -79.183627 | 05 Oct 2014 | QCAZ:A:58308 | RDT0088 | 19.55  0.6 | ND |
| Dendrobatidae | *Epipedobates currulao* | 291.941 | 99 | La Barra, Valle del Cauca, CO:  3.985064, -77.376723 | 26 Nov 2014 | ANDES:A:2455 | RDT0147 | 17.19  0.7 | F |
| Dendrobatidae | *Epipedobates currulao* | 352.296 | 105 | Ladrilleros, Valle de Cauca, CO:  3.950958, -77.358312 | 27 Nov 2014 | ANDES:A:2459 | RDT0151 | 17.62  0.4 | ND |
| Dendrobatidae | *Epipedobates boulengeri* | 262.467 | 94 | Maragrícola, Nariño, CO:  1.68084, -78.74924 | 09 Dec 2014 | ANDES:A:2468 | RDT0173 | 18.30  0.6 | F |
| Dendrobatidae | *Epipedobates boulengeri* | 157.450 | 77 | Maragrícola, Nariño, CO:  1.68084, -78.74924 | 09 Dec 2014 | ANDES:A:2469 | RDT0174 | 18.68  0.6 | F |
| Dendrobatidae | *Hyloxalus awa* | 1.048 | 7 | Otongachi, Santo Domingo, EC:  -0.321902, -78.952072 | 17 Sep 2014 | QCAZ:A:58267 | RDT0047 | 22.24  1.0 | ND |
| Dendrobatidae | *Hyloxalus awa* | 0.168 | 4 | Otongachi, Santo Domingo, EC:  -0.321902, -78.952072 | 17 Sep 2014 | QCAZ:A:58268 | RDT0048 | 22.26  1.0 | ND |
| Dendrobatidae | *Hyloxalus awa* | 3.002 | 12 | Otongachi, Santo Domingo, EC:  -0.321902, -78.952072 | 17 Sep 2014 | QCAZ:A:58269 | RDT0049 | 22.79  1.3 | ND |
| Dendrobatidae | *Hyloxalus awa* | 0.787 | 1 | Valle Hermoso, Santo Domingo, EC:  -0.086217, -79.272098 | 15 Sep 2014 | QCAZ:A:58241 | RDT0021 | 18.58  0.7 | ND |
| Dendrobatidae | *Hyloxalus awa* | 0.778 | 1 | Valle Hermoso, Santo Domingo, EC:  -0.086217, -79.272098 | 15 Sep 2014 | QCAZ:A:58242 | RDT0022 | 19.98  1.0 | ND |
| Dendrobatidae | *Hyloxalus awa* | 9.309 | 3 | Valle Hermoso, Santo Domingo, EC:  -0.086217, -79.272098 | 15 Sep 2014 | QCAZ:A:58243 | RDT0023 | 17.80  0.8 | ND |
| Dendrobatidae | *Hyloxalus awa* | 0.000 | 0 | Valle Hermoso, Santo Domingo, EC:  -0.086217, -79.272098 | 16 Sep 2014 | QCAZ:A:58244 | RDT0024 | 18.23  0.7 | ND |
| Dendrobatidae | *Hyloxalus shuar* | 3.010 | 5 | Punguinzta, Zamora-Chinchipe, EC:  -3.950288, -78.887773 | 29 Sep 2014 | QCAZ:A:58289 | RDT0069 | 20.79  0.8 | ND |
| Dendrobatidae | *Hyloxalus* sp. Agua Azul | 1.630 | 8 | Santa Maria, Boyacá, CO: 4.84756,  -73.271079 | 03 Nov 2014 | ANDES:A:2426 | RDT0118 | 15.1  0.4 | ND |
| Dendrobatidae | *Hyloxalus toachi* | 0.000 | 0 | Lita, Esmeraldas, EC:  0.85401, -78.463594 | 11 Sep 2014 | QCAZ:A:58229 | RDT0009 | 16  0.4 | ND |
| Dendrobatidae | *Hyloxalus toachi* | 0.000 | 0 | Lita, Esmeraldas, EC:  0.85401, -78.463594 | 11 Sep 2014 | QCAZ:A:58230 | RDT0010 | 21.00  NA | F |
| Dendrobatidae | *Leucostethus fugax* | 0.287 | 3 | Canelos, Pastaza, EC:  -1.584533, -77.794459 | 08 Oct 2014 | QCAZ:A:58311 | RDT0091 | 19.42  0.7 | ND |
| Dendrobatidae | *Leucostethus fugax* | 1.027 | 3 | Canelos, Pastaza, EC:  -1.584533, -77.794459 | 08 Oct 2014 | QCAZ:A:58312 | RDT0092 | 16.15  0.5 | ND |
| Dendrobatidae | *Leucostethus fugax* | 2.259 | 5 | Canelos, Pastaza, EC:  -1.584533, -77.794459 | 10 Oct 2014 | QCAZ:A:58315 | RDT0095 | 19.49  0.8 | ND |
| Dendrobatidae | *Leucostethus fugax* | 1.249 | 7 | Canelos, Pastaza, EC:  -1.584533, -77.794459 | 10 Oct 2014 | QCAZ:A:58316 | RDT0096 | 19.07  0.7 | ND |
| Dendrobatidae | *Leucostethus fugax* | 4.568 | 8 | Canelos, Pastaza, EC:  -1.584533, -77.794459 | 10 Oct 2014 | QCAZ:A:58317 | RDT0097 | 17.42  0.6 | ND |
| Dendrobatidae | *Leucostethus fugax* | 0.413 | 3 | Canelos, Pastaza, EC:  -1.584533, -77.794459 | 10 Oct 2014 | QCAZ:A:58318 | RDT0098 | 16.07  0.5 | ND |
| Dendrobatidae | *Leucostethus fugax* | 3.165 | 7 | Canelos, Pastaza, EC:  -1.584533, -77.794459 | 10 Oct 2014 | QCAZ:A:58319 | RDT0099 | 19.9  1.0 | F |
| Dendrobatidae | *Leucostethus fugax* | 1.161 | 4 | Canelos, Pastaza, EC:  -1.584533, -77.794459 | 10 Oct 2014 | QCAZ:A:58320 | RDT0100 | 17.81  0.6 | ND |
| Dendrobatidae | *Oophaga sylvatica* | 61,743.240 | 152 | El Pailón, Nariño, CO:  1.410098, -78.238718 | 13 Dec 2014 | ANDES:A:2477 | RDT0182 | 35.31  2.8 | ND |
| Dendrobatidae | *Oophaga sylvatica* | 8,506.890 | 159 | Villa Hermosa, Santo Domingo, EC:  -0.268356, -79.12051 | 17 Sep 2014 | QCAZ:A:58255 | RDT0035 | 29.46  1.9 | ND |
| Dendrobatidae | *Oophaga sylvatica* | 24,691.799 | 189 | Villa Hermosa, Santo Domingo, EC:  -0.268356, -79.12051 | 17 Sep 2014 | QCAZ:A:58256 | RDT0036 | 29.98  1.7 | ND |
| Dendrobatidae | *Oophaga sylvatica* | 20,913.042 | 181 | Villa Hermosa, Santo Domingo, EC:  -0.268356, -79.12051 | 17 Sep 2014 | QCAZ:A:58257 | RDT0037 | 26.17  1.4 | ND |
| Dendrobatidae | *Oophaga sylvatica* | 14,406.872 | 175 | Villa Hermosa, Santo Domingo, EC:  -0.268356, -79.12051 | 17 Sep 2014 | QCAZ:A:58258 | RDT0038 | 27.57  1.8 | ND |
| Dendrobatidae | *Phyllobates aurotaenia* | 96.580 | 54 | La Barra, Valle del Cauca, CO: 3.985532, -77.376676 | 26 Nov 2014 | ANDES:A:2456 | RDT0148 | 23.11  1.6 | ND |
| Dendrobatidae | *Phyllobates aurotaenia* | 49.595 | 48 | Ladrilleros, Valle de Cauca, CO: 3.951089, -77.358173 | 27 Nov 2014 | ANDES:A:2457 | RDT0149 | 27.73  1.9 | F |
| Dendrobatidae | *Phyllobates aurotaenia* | 261.920 | 81 | Termales, Chocó, CO:  5.620467, -77.420829 | 17 Nov 2014 | ANDES:A:2434 | RDT0126 | 21.19  0.8 | ND |
| Dendrobatidae | *Phyllobates aurotaenia* | 1,421.973 | 118 | Termales, Chocó, CO:  5.620467, -77.420829 | 17 Nov 2014 | ANDES:A:2435 | RDT0127 | 20.91  0.8 | ND |
| Dendrobatidae | *Rheobates palmatus* | 1.525 | 4 | Las Brisas, Cundinamarca, CO: 4.432886, -73.920631 | 09 Nov 2014 | ANDES:A:2430 | RDT0122 | 29.41  2.89 | F |
| Dendrobatidae | *Rheobates palmatus* | 1.502 | 2 | Las Brisas, Cundinamarca, CO: 4.432886, -73.920631 | 09 Nov 2014 | ANDES:A:2431 | RDT0123 | 27.75  2.35 | F |
| Dendrobatidae | *Rheobates palmatus* | 0.476 | 1 | Las Brisas, Cundinamarca, CO: 4.432886, -73.920631 | 09 Nov 2014 | ANDES:A:2432 | RDT0124 | 29.66  3.21 | F |
| Dendrobatidae | *Rheobates palmatus* | 0.961 | 1 | Las Brisas, Cundinamarca, CO: 4.432886, -73.920631 | 09 Nov 2014 | ANDES:A:2433 | RDT0125 | 28.77  3.01 | F |
| Dendrobatidae | *Silverstoneia* aff*. gutturalis* | 2.156 | 3 | El Valle, Chocó, CO:  6.122592, -77.44447 | 18 Nov 2014 | ANDES:A:2439 | RDT0131 | 17.14  0.35 | ND |
| Dendrobatidae | *Silverstoneia* aff*. gutturalis* | 12.848 | 3 | El Valle, Chocó, CO:  6.122592, -77.44447 | 18 Nov 2014 | ANDES:A:2440 | RDT0132 | 19.11  0.5 | ND |
| Dendrobatidae | *Silverstoneia* aff*. gutturalis* | 3.374 | 9 | El Valle, Chocó, CO:  6.122592, -77.44447 | 18 Nov 2014 | ANDES:A:2441 | RDT0133 | 18.1  0.55 | F |
| Dendrobatidae | *Silverstoneia* aff*. gutturalis* | 4.867 | 10 | El Valle, Chocó, CO:  6.122592, -77.44447 | 18 Nov 2014 | ANDES:A:2442 | RDT0134 | 17.8  0.25 | M |
| Dendrobatidae | *Silverstoneia* aff*. gutturalis* | 0.339 | 1 | El Valle, Chocó, CO:  6.122592, -77.44447 | 18 Nov 2014 | ANDES:A:2443 | RDT0135 | 18.4  0.55 | F |
| Dendrobatidae | *Silverstoneia* aff*. gutturalis* | 33.439 | 4 | El Valle, Chocó, CO:  6.122592, -77.44447 | 18 Nov 2014 | ANDES:A:2444 | RDT0136 | 18.18  0.6 | F |
| Dendrobatidae | *Silverstoneia* aff*. gutturalis* | 0.134 | 3 | El Valle, Chocó, CO:  6.122592, -77.44447 | 18 Nov 2014 | ANDES:A:2445 | RDT0137 | 15.31  0.35 | M |
| Dendrobatidae | *Silverstoneia* aff*. gutturalis* | 10.730 | 3 | El Valle, Chocó, CO:  6.122592, -77.44447 | 18 Nov 2014 | ANDES:A:2446 | RDT0138 | 18.24  0.6 | F |
| Dendrobatidae | *Silverstoneia* aff*. gutturalis* | 5.130 | 4 | El Valle, Chocó, CO:  6.122592, -77.44447 | 18 Nov 2014 | ANDES:A:2447 | RDT0139 | 17.63  0.4 | ND |
| Dendrobatidae | *Silverstoneia erasmios* | 9.949 | 15 | Los Tubos, Valle de Cauca, CO: 3.847596, -76.788457 | 28 Nov 2014 | ANDES:A:2466 | RDT0158 | 17.74  0.5 | ND |
| Dendrobatidae | *Silverstoneia erasmios* | 2.429 | 15 | Los Tubos, Valle de Cauca, CO: 3.847596, -76.788457 | 28 Nov 2014 | ANDES:A:2467 | RDT0159 | 17.23  0.4 | ND |
| Leptodactylidae | *Lithodytes lineatus* | 0.000 | 0 | Zumbi, Zamora-Chinchipe, EC:  -3.888602, -78.786518 | 28 Sep 2014 | QCAZ:A:58286 | RDT0066 | 19.56  0.65 | ND |
| Leptodactylidae | *Lithodytes lineatus* | 0.000 | 0 | Zumbi, Zamora-Chinchipe, EC:  -3.888602, -78.786518 | 28 Sep 2014 | QCAZ:A:58287 | RDT0067 | 19.11  0.6 | ND |
