## Supplementary Table S6 for "Passive accumulation of alkaloids in inconspicuously colored frogs refines the evolutionary paradigm of acquired chemical defenses"

**Table S6**. A list of classes (inferred by NPClassifier) and specific classes (inferred by ClassyFire) of metabolites assigned to the alkaloid pathway detected in skins of *Silverstoneia flotator* (Dendrobatidae). Where these compounds are also found in *Eleutherodactylus cystignathoides* (Eleutherodactylidae) is indicated in the fourth column. In some cases, two or more compounds with identical specific-class annotations may be assigned to different classes (e.g., biguanides assigned to “Acyclic guanidine alkaloids” and “Polyamines”). The fifth column lists if the compound class or specific class fall under the broad categories of lipophilic alkaloids listed in the Daly database and the sixth column designates whether or not the molecular formula and monoisotopic mass are found in the Daly database.

| **Class (inferred by NPClassifier)** | **Specific Class (inferred by ClassyFire)** | **Number in *S. flotator*** | **Number in *El. cystignathoides*** | **Database alkaloid class?** | **Database molecular formula?** |
| --- | --- | --- | --- | --- | --- |
| Acyclic guanidine alkaloids | Biguanides | 1 | 0 | No | No |
| Acyclic guanidine alkaloids | Guanidines | 1 | 0 | No | No |
| Carboline alkaloids | Amino acids and derivatives | 1 | 1 | No | No |
| Isoindole alkaloids | Benzenoids | 1 | 0 | No | No |
| Isoindole alkaloids | Phthalimides | 1 | 1 | No | No |
| Isoquinoline alkaloids | Aralkylamines | 1 | 1 | No | No |
| Isoquinoline alkaloids | Diazanaphthalenes | 1 | 0 | No | No |
| Phenazine alkaloids | Quinoxalines | 1 | 1 | No | No |
| Phenylalanine-derived alkaloids | Alkyl-phenylketones | 1 | 1 | No | No |
| Phenylethylamines | 1-hydroxy-2-unsubstituted benzenoids | 1 | 0 | No | No |
| Phenylethylamines | Methoxyphenols | 1 | 0 | No | No |
| Piperidine alkaloids | Benzylpiperidines | 1 | 1 | Yes | No |
| Polyamines | Azacyclic compounds | 1 | 1 | No | No |
| Polyamines | Amino acids and derivatives | 1 | 1 | No | No |
| Polyamines | Aminopyrimidines and derivatives | 1 | 1 | No | No |
| Polyamines | Biguanides | 2 | 1 | No | No |
| Polyamines | Biopterins and derivatives | 2 | 2 | No | No |
| Polyamines | Diazines | 1 | 1 | No | No |
| Polyamines | Heteroaromatic compounds | 1 | 1 | No | Yes |
| Polyamines | N-acyl amines | 1 | 1 | No | No |
| Polyamines | Tetrazoles | 2 | 0 | No | No |
| Polyamines | Trialkylamines | 1 | 0 | No | No |
| Purine alkaloids | 6-oxopurines | 5 | 4 | No | No |
| Purine alkaloids | Guanidines | 1 | 1 | No | No |
| Purine alkaloids | N-arylpiperazines | 1 | 0 | No | No |
| Purine alkaloids | Purines and purine derivatives | 1 | 1 | No | No |
| Purine alkaloids | Purinones | 1 | 0 | No | No |
| Purine alkaloids | Xanthines | 2 | 2 | No | No |
| Pyridine alkaloids | Pyridinecarboxamides | 1 | 1 | Yes | No |
| Pyridine alkaloids | Thiadiazoles | 1 | 1 | Yes | No |
| Pteridine alkaloids | 6-oxopurines | 2 | 0 | No | No |
| Pteridine alkaloids | Amino acids | 2 | 1 | No | No |
| Pteridine alkaloids | Amino acids and derivatives | 4 | 3 | No | No |
| Pteridine alkaloids | Anisoles | 1 | 1 | No | No |
| Pteridine alkaloids | Flavins | 1 | 1 | No | No |
| Pteridine alkaloids | N-acetylarylamines | 3 | 0 | No | No |
| Pteridine alkaloids | Pteridines and derivatives | 1 | 0 | No | No |
| Pteridine alkaloids | Pterins and derivatives | 6 | 4 | No | No |
| Pteridine alkaloids | Thiadiazoles | 1 | 1 | No | No |
| Pyrazine and piperazine alkaloids | Quinoxalines | 1 | 0 | No | No |
| Quinazoline alkaloids | Quinoxalines | 1 | 0 | No | No |
| Quinazoline alkaloids | Benzodiazines | 1 | 1 | No | No |
| Quinolizidine alkaloids | Quinolizidines | 1 | 0 | Yes | Yes |
| Simple amide alkaloids | N,N-dialkyl-m-toluamides | 1 | 1 | No | Yes |
| Simple indole alkaloids | Benzenesulfonamides | 1 | 0 | No | No |
| Steroidal alkaloids | Amino acids and derivatives | 1 | 1 | No | No |
| Tropane alkaloids | Epibatidine analogues | 1 | 0 | Yes | Yes |
